# Tandem electrospinning for heterogeneous nanofiber patterns

**DOI:** 10.1101/830950

**Authors:** Paul Wieringa, Roman Truckenmuller, Silvestro Micera, Richard van Wezel, Lorenzo Moroni

## Abstract

Smart nanofibrillar constructs can be a promising technological solution for many emerging and established fields, facilitating the potential impact of nano-scale strategies to address relevant technological challenges. As a fabrication technique, electrospinning (ESP) is relatively well-known, accessible, economic, and fast, though until now has shown limitation over control and design of the fibrillar constructs which can be produced. Here, we introduce “Tandem Electrospinning” (T-ESP), a novel technique able to create increasingly complex patterns of fibrous polymeric constructs on a micro and nano-scale. Modifying a standard ESP configuration results in a focusing electric field that is able to spatially define the deposition pattern of multiple polymer jets simultaneously. Additional spatially defined heterogeneity is achieved by tuning polymer solution properties to obtain a gradient of fiber alignment. Heterogeneous fibrous meshes are created with either random, aligned, or a divergent fiber patterns. This approach holds potential for many fields, with application examples shown for Tissue Engineering and Separation Technologies. The technique outlined here provides a rapid, versatile, and accessible method for polymeric nanofabrication of patterned heterogeneous fibrous constructs. Polymer properties are also shown to dictate fiber alignment, providing further insight into the mechanisms involved in the electrospinning fabrication process.

## Introduction

Recent years have seen a renewed interest in electrospinning (ESP) for micro- and nano-scale fabrication, driven by new applications in the fields of environment and security, such as filtration or particle sensing, biomedical technologies, in form of tissue scaffolds or drug release platforms, and in the energy sector, with potential for improving both solar and fuel cell technologies^1–6^. The promise of creating micro- and nano-sized fibers in a simple, relatively inexpensive and high-throughput manner has spurred development of ESP fabrication techniques, leading to the production of fibrous meshes with high surface area and tunable porosity.

Conventional ESP results in a chaotic fibrous mesh, narrowing the application of this form of ESP to areas such as textile and filtration industries. Strategies have been developed to produce fiber constructs useful for other applications, but these approaches still exhibit limitations in terms of spatially defined fiber deposition and orientation, and typically produce simple polymer constructs consisting of a single fiber type. ESP across a planar target gap electrode or patterned target electrodes produces simple parallel fibers or cross-hatched fiber patterns, respectively, with limited spatial definition and restricted to single fiber composition or layer-by-layer fiber assemblies^1, 7–11^. In general, deposition and spatial control of multiple polymer fibers is hampered by the mutual repulsion of multiple charged jets^12–14^. Attempts to create multimodal fiber constructs have relied on the chaotic fluctuations of the ‘whipping’ polymer jets to achieve overlap between fiber populations, intrinsically precluding any form of fiber orientation. ESP onto a rotating mandrel also produces simple parallel fibers, with increased complexity possible by simultaneously electrospinning different polymers onto a rotating mandrel^15^. However, more elaborate, spatially defined multimodal patterns are not possible. The most detailed fiber patterns have been created by near-field electrospining (NFES) or melt electrospinning, which are less accessible and unable to simultaneously deposit multiple fiber types within a confined region^16, 17^.

Here, we describe the development of T-ESP techniques, which allow for simultaneous electrospinning of up to three polymer fibers. Using modifications to a traditional ESP setup, we can achieve a high degree of control of planar ESP patterning. Control over multiple jets is achieved by manipulating the electrostatic field, resulting in the creation of ESP multi-material fibrous constructs. This allows ESP fiber patterns of intermingled, heterogeneous fiber populations in random, parallel or divergent (“Y-shaped”) orientations. Finally, we show how this facile and economic fabrication method can be used to mimic branched structures found in biological systems and present an example of a divergent tissue scaffold supporting spatially defined differences in neurite outgrowth.

### Experimental Section

#### Polymer Solution Preparation and Characterization

All polymers were dissolved in associated solvent solutions overnight (See Supplementary Table S1 for complete solution details) and appropriate dyes were added at 0.01% w/v. NaCl (Sigma Aldrich GmbH, Germany) was added to HW and HW*_mv_* polymer solutions at final concentration of 0.05 mg/ml. Conductivity was measured using a custom gold parallel plate apparatus at 20 °C (see Supplementary Material S12 for full description) and viscosity was measured using a Brookfield DV-E with an s21 mandrel at 30 °C and 100 rpm.

#### Electrospinning for T-ESP Development

A custom ESP setup was used with an environmental chamber (25°C, 30% humidity) and a large upper parallel plate (30 cm by 20 cm). ESP was performed at 25 KV, a working distance of 20 cm, a flow rate of 1 ml/hr and for an ESP interval of 1 minute. For tandem ESP and triple ESP, needles were mounted along the Y axis of the collector target (Figure 1B) at an inter-needle separation of 1 cm and 5 cm, respectively. A 14 mm coverslip (Menzel-Glaser) was the target substrate used to collect deposited fibers, placed on the collector electrodes of either a parallel or epsilon arrangement (see Supplementary Figure S8).

**Figure 1.**
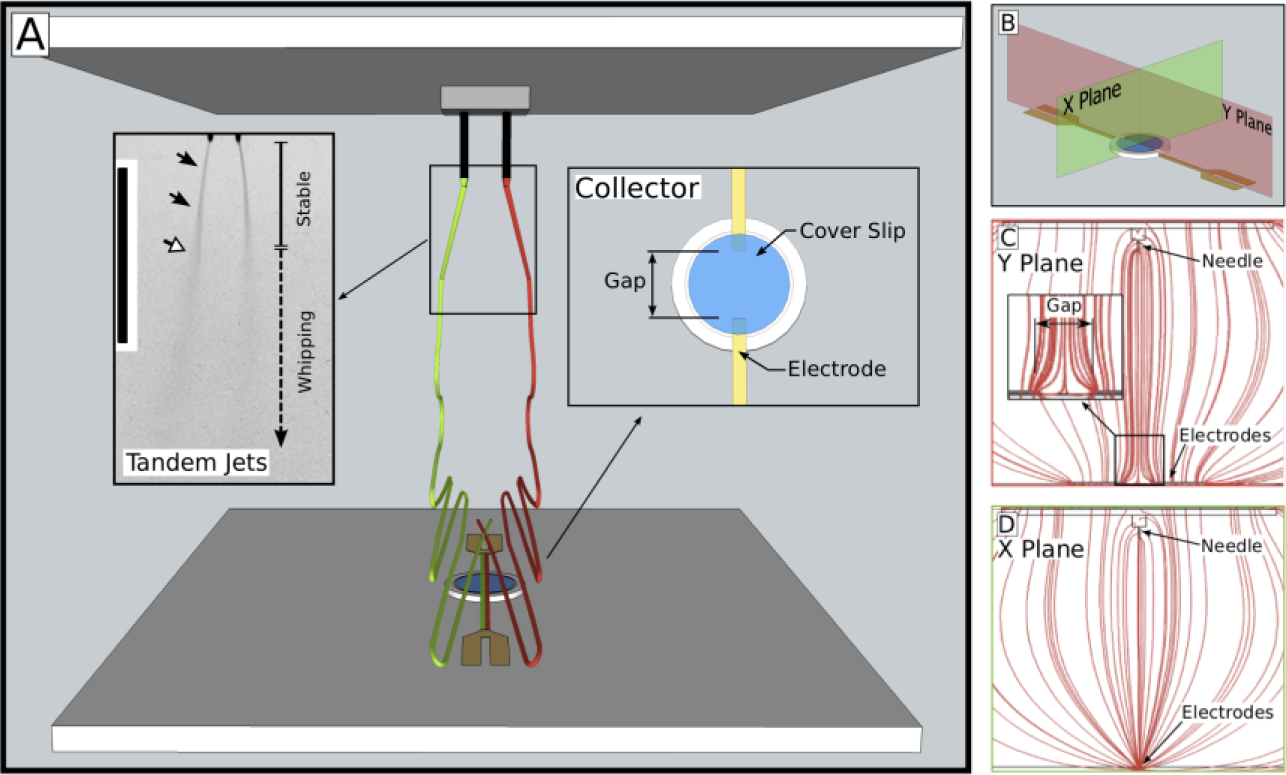
The Tandem ESP Apparatus. (A) A depiction showing an upper parallel plate to which 2 needles are mounted and high voltage is applied and a base plate with a gap electrode arrangement (detail shown in Inset: “Collector”). A 10 mm gap was made between two 2 mm wide electrodes. Shown is the placement of a 14 mm glass coverslip on top of the two electrodes, used to collect fibers patterned across the gap. ESP with single needles were centered over the coverslip. Tandem and triple ESP needles were arranged along the X axis (see C for orientation). Also shown is a representation of two simultaneous jets (red and green) as well as an image of the polymer jets of two LW polymer solutions (Inset: “Tandem Jets”; scale bar: 5 cm); dark arrows indicate the stable jet phase and a white arrow indicates the onset of the whipping phase, where the jet becomes too small to be easily visible. (B) The gap electrode configuration was modeled to show the electric field generation in both the X and Y plane. (C) The electric field in the Y plane of the gap electrode configuration shows the divergent electric field (inset) which causes fibers to align across the gap. (D) The same electric field in the orthogonal X plane clearly shows the focusing electric field this electrode arrangement creates.

#### Electrospinning T-ESP Scaffolds for Specific Applications

Cell scaffolds were prepared using the epsilon electrode configuration shown in S9 and fibers were deposited on flexible mesh rings (outer diameter 15 mm, inner diameter 12 mm) in lieu of coverslips for improved handling. A working distance of 20 cm and a voltage of 20 KV was used for an ESP time of 30 seconds. To ensure jet stability, the LW solution had a flow rate of 1 ml/hr and a flow rate of 0.5 ml/hr for the LW-collagen solution. Parallel T-ESP scaffolds for liquid phase separation were prepared with the standard gap electrode configuration using an LW solution (1.0 ml/hr) and a 10% PVA solution (0.5 ml/hr) on glass coverslips. Scaffold were immersed in methanol with 5% paraformaldehyde for 24 hrs to crosslink the PVA fibers and then air dried for 24 hrs before use.

#### Analysis of ESP Process

Images of whipping were captured by a Luminix DMC G3 and fiber pattern images were stitched together using a Nikon Eclipse Ti with an automated stage at 10x magnification. Fibers were gold sputter-coated and examined with a XL 30 ESEM-FEG (Phillips) operating at 10 kV. A minimum of 100 fibers were measured per population and minimum of 5 images per fiber population were evaluated for fiber orientation using the OrientationJ plugin^45^, providing a coherence value between 0 (isotropic) and 1 (perfectly anisotropic). Fiber orientation was also evaluated by creating an FFT image of SEM images^46^.

#### Cell Culture

Dorsal root ranglia (DRGs) were explanted from 2 day old rat pups (Wistar Unilever: HsdCpb:WA). All procedures followed national and European laws and guidelines and were approved by the local ethical committee. Briefly, rats were sacrificed by cervical dislocation under general anaesthesia (4% Isoflurane) and then decapitated. Individual ganglia were removed from the spinal column and nerve roots were stripped under aseptic conditions with the aid of a stereomicroscope. DRGs were cut in half to expose interior neurites and placed at the divergent junction of the tandem ESP scaffold. DRGs were cultures in NeuralBasal A-medium with B27 supplement, 0.5 mM L-glutamine and pen/strep added (Gibco/Invitrogen) and 10 ng/ml NGF (Sigma Aldrich). Cultures were maintained for 5 days, with medium refreshed after Day 1 and Day 3. Cells were fixed with ice cold 2% paraformaldehyde (PFA) for 15 minutes at 4°C, then permeabilized for 10 minutes with 0.1% TritonX-100. Cultures were blocked for 1 hour in 5% Normal Goat Serum and 1% BSA in a TRIS buffered solution (TBS), followed by a 16 hr incubation with a mouse anti-b3-tubulin primary antibody (1:1000,Abcam), a triple wash in 1% BSA TBS solution and a 16 hr incubation with an Alexa 546 anti-mouse secondary antibody (1:1000, Invitrogen) with 1% Normal Goat serum. Scaffolds were then washed and mounted with Mowiol 4-88 with 2.5% DABCO.

*Additional methodology information is available in Supplementary Material section*.

## Results and Discussion

### Electrode configuration focuses multiple polymer jets for controlled overlapping fiber deposition

Traditional ESP process can be described as an electrostatic extrusion of a charged polymer solution from a supply needle towards a grounded collector. During this process, a polymer jet is formed which is initially straight but then enters an unstable whipping phase. This chaotic motion further extrudes the polymer jet and accelerates solvent evaporation to form solid polymer fibers^18^. To achieve more control over fiber deposition, this study manipulated the electrostatic field by using a grounded target comprised of millimeter-wide electrodes with a centimeter gap (Figure 1). The electric field across the gap is known to exert forces on the fibers during the whipping phase (Figure 1A), leading to fiber alignment across the gap^19^. However, the use of millimeter wide electrodes in this configuration (Figure 1C, 1A Inset: “Collector”) also achieved a focusing effect of the electric field in the orthogonal plane (Figure 1D) to further direct the evolution of the whipping phase.

ESP of one polymer solution with this arrangement led to the spatially confined deposition of aligned fibers across this gap (Figure 2A). Yet, this novel electrode architecture produced an electric field able to confine the simultaneous deposition of two ESP jets to produced aligned parallel deposition of different fiber populations (Figure 2B). To the best of our knowledge, this is the first report of concurrent and controlled deposition of different populations of aligned fibers with a well-defined region of overlap (Figure 2C).

**Figure 2.**
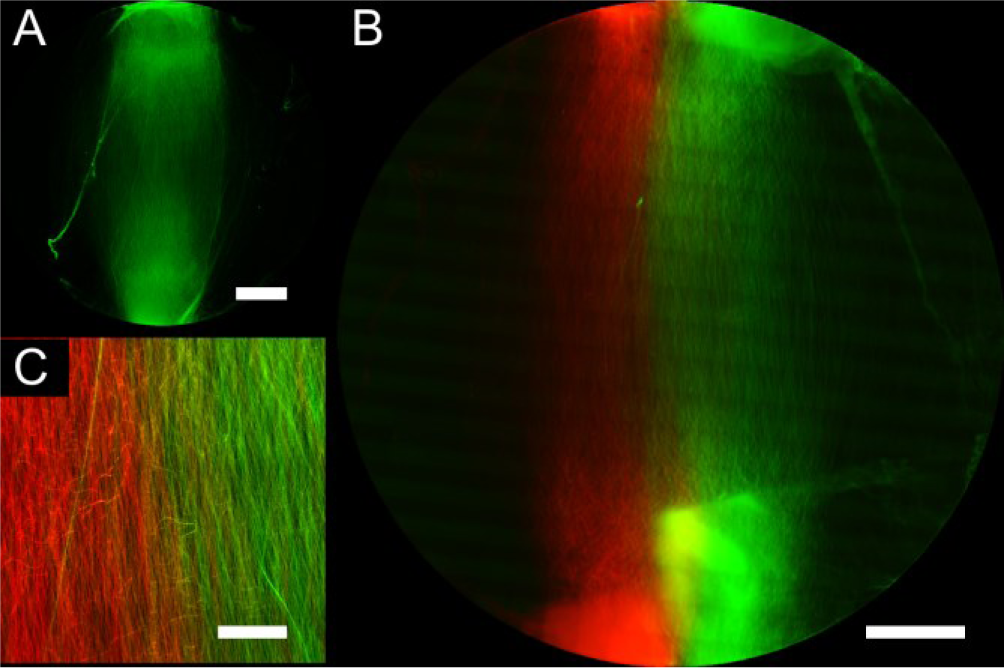
Aligned Tandem ESP constructs. (A) Aligned ESP of a single jet of LW polymer solution, confined to the approximate width of the electrodes (scale bar: 2 mm). Fibers have a diameter of 0.756 ± 0.160 μm and are well aligned, with a coherence measurement of 0.780 ± 0.059 (1 is perfectly aligned, 0 is isotropic; see Figure S1 for additional information). (B) The tandem ESP deposition pattern of two identical LW polymer solutions with Macrolex® Red G and Green dyes added, respectively (scale bar: 2 mm). This particular polymer solution produces consistent electrospun fibers (green: 0.653±0.143 μm; red: 0.637±0.136 μm) and achieves a high degree of alignment when using a gap target electrode configuration (green: 0.734±0.115; red:0.817±0.097). (C) A magnified view of the overlapping regions between the two fiber populations is shown, clearly depicting the defined yellow gradient transition from red to green (scale bar: 2 mm).

Despite initial mutual repulsion of the two jets during the stable jet phase (Figure 1A Inset: “Tandem Jets”), the focused electric field provided sufficient force upon the fibers during the whipping phase to cause colocalized deposition. Fluorescence microscopy (Figure 2C) revealed that the two regions were overlapping with no distinct boundary region between fiber populations. The fibers showed a relatively homogenous diameter and were well aligned, with no statistically significant differences between populations.

### Polymer properties dictate alignment for gap electrode ESP

The degree of fiber alignment achieved was found to be dependent on the polymer used. This provided the possibility of further control over fiber deposition and degree of multimodal complexity achievable. The polymer used in the experiment above (designated LW) is of a family of segmented block copolymers compromised of poly(ethylene oxide terephthalate) (PEOT) and poly(butylene terephthalate) (PBT), synthesized using PEOT and PBT segments of specific lengths added in known proportions to produce a randomly distributed block copolymer. Originally devised as materials for tissue scaffold fabrication and controlled drug delivery systems, the polymer properties are tuned per application by controlling the length of the initial PEOT segments and varying proportions of PEOT to PBT^20–22^. A polymer with initial PEOT segments of 300 Da and a PEOT/PBT of 55/45 has the designation of 300PEOT55PBT45.

The subset of polymers used in this study was selected for having similar molecular weights of approximately 50 kDa but having different compositions and distinct mechanical properties. A distinguishing characteristic of the different polymers selected was the entanglement molecular weight (M*_e_*); this is a measure of the chain length between physical entanglement points of polymer chains in a bulk polymer and has an impact on the viscoelastic properties of the polymer^23^. M*_e_*, along with viscosity and solution conductivity, has been cited as an important parameter in describing the polymer solutions used for ESP^24–26^. The three polymers selected were: 300PEOT55PBT45 (M*_e_*: 250 Da) and designated in this study as LW (low M*_e_*); 300PEOT70PBT30 (M*_e_*: 450 Da), referred to as IW (intermediate M*_e_*); and 1000PEOT70PBT30 (M*_e_* : 710 Da), called here HW (high M*_e_*)^27^. These were prepared as polymer solutions in solvent blends of chloroform and hexafluoroisopropanol (HFIP; see Supplementary Material, Table S1 and S2 for a details on polymers and solutions used).

Despite similar mean molecular weights and a 20% w/v polymer concentration, the LW, IW and HW polymers solutions demonstrated different viscosities of 120, 125 and 388 cP which produced fiber diameters of 0.86±0.28 0.74±0.34, and 1.03±0.35 □m respectively. Viscosity of a polymer solution is dependent on polymer chain entanglements, or overlap, with a higher polymer concentration or molecular weight increasing the availability of polymer chains to entangle in solution and leading to an observable increase in solution viscosity; this is also dependent on the solubility of the polymer, as evidenced here by the higher viscosity of the HW solution^28^. Viscosity is often correlated to either changes in fiber size or quality, exemplified in the current study by the higher viscosity HW solution which produced larger fibers^24, 29, 30^.

Unexpectedly, image analysis of fiber orientation found statistically significant decrease (p < 0.01) in alignment with increasing values of M*_e_*. Using a score from 0 to 1 (random to aligned), alignments of 0.773±0.089, 0.641±0.102, and 0.33±0.077 were measured for LW, IW, and HW fibers, respectively. To account for differences in viscosity, the concentration of the HW polymer solution was adjusted to achieve a similar viscosity of 122 cP (referred to as HW*_mv_*, ‘matched viscosity’; see Table S2). However, the resulting fiber alignment of 0.222±0.0698 indicated that viscosity, thus entanglement in the initial polymer solution, was not an effective predictor of fiber alignment. Instead, M*_e_* as a bulk polymer property appeared to be the predominant factor. Until now, M*_e_* has been used to describe the viscoelasticity of the polymer solution as it relates to the formation of ‘beads-on-a string’ fiber morphology formed in the initial stages of the polymer jet^24^. This finding suggested a potential role of M*_e_* on fiber morphology after they have returned to a bulk polymer state later in the whipping phase.

### M*_e_* affects whipping phase evolution and fiber alignment

The difference between whipping phases of the different polymers is clearly observed from the macroscopic patterns of fiber deposition, with the size and shape of the region of deposition dependent on how the whipping phase develops. The highly entangled LW polymer deposits fibers along the length of the electrodes, indicating a smooth, elongated progression of the whipping phase. In comparison, the HW polymer has fewer fibers along the length of the electrodes and a wider, more circular deposition in the electrode gap, suggesting a more abrupt, arrested evolution of the whipping fiber (Supplementary Material Figure S1). It is suggested here that M*_e_* influences both fiber formation during the whipping phase and the overall evolution of the whipping phase, accounting for many of the observed differences between the polymers fibers.

In addition to the initial perturbations (bending) preceding the onset of the whipping instability, it has been previously reported that the fiber can also experience secondary and tertiary bending during the whipping phase. The consequence is that fibers are not necessarily smooth and straight, but can have a wavy, corrugated appearance^18^. Considering also the strain of the solidifying fiber during the whipping phase and the correlation between degree of entanglement and plastic deformation, it is proposed here that a fiber with a higher entanglement molecular weight (M*_e_*) experiences more plastic deformation during the whipping phase^31, 32^. In turn, this increased drawing results in a curled, corrugated fiber, likely caused by non-axisymmetric residual stress^33^. In comparison, a fiber of a highly entangled polymer would have increased elasticity, resisting deformation, and would incur less residual stress to produce a straight fiber morphology. It should also be remarked that the long working distance used (20 cm) excludes the curled fibers to be the result of buckling^34^.

We suggest that the electric field imposes orientation along the length of the fibers, such that a general ‘global’ scale alignment is achieved and fiber orientation is not entirely random. However, the resulting microscopic alignment of the fiber also depends on how corrugated the fiber has become. As observed, higher M*_e_* polymers would produce fibers with increasing disorder, although not entirely random. Since strain and disentanglement can also be affected by differences in molecular weight, use of these particular polymers proved essential in identifying the role of M*_e_*^35^.

These additional perturbations may also disrupt the whipping profile development, explaining the observed differences in the elongated deposition of fibers. An elongated whipping profile may also increase the degree of microscopic fiber alignment of the LW polymer fibers by improving global fiber alignment and effectively stretching out fiber perturbations.

### Heterogeneous fiber patterns can be tuned by adjusting polymer solution properties

T-ESP was extended to produce heterogeneous fiber populations, using the LW and the HW polymers as far extremes offered by this family of polymers in terms of both fiber size and fiber alignment. Heterogeneous T-ESP produced the expected regions of aligned LW fibers (0.699±0.104), less aligned HW fibers (0.401±0.06), and an overlapping region between the two populations. Unexpectedly, this also resulted in a fiber deposition pattern with an extreme spatial bias, with the HW fibers now shifted to the edge of the deposition region (Figure 3A).

**Figure 3.**
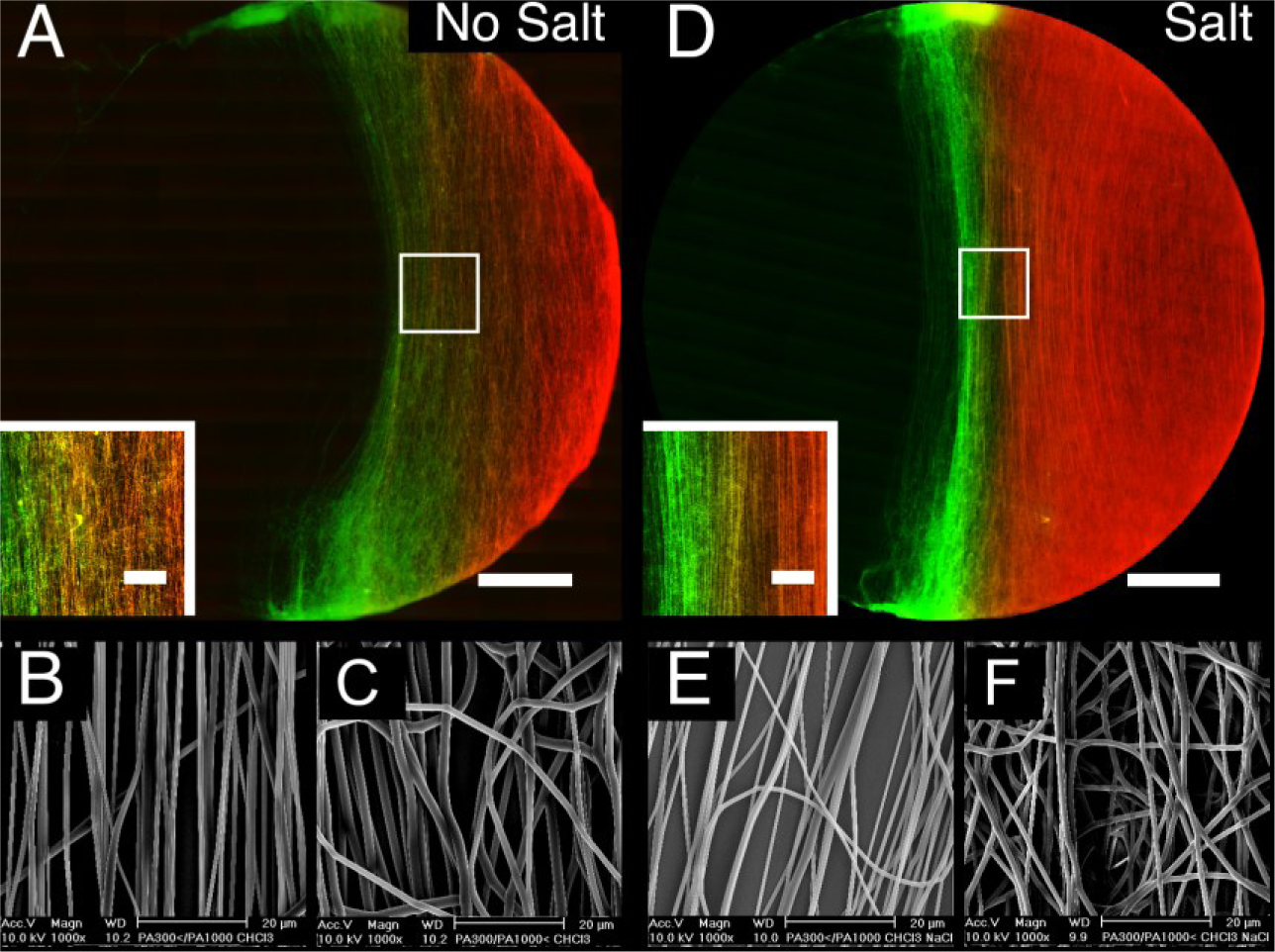
Heterogeneous Tandem ESP Scaffolds. (A) Tandem ESP fibers of LW (green) and HW (red) produced a biased pattern of fiber deposition, though an overlapping gradient region between the two fiber populations was still maintained (inset). (B) The LW fibers had a diameter of 0.723 ± 0.261 μm and were well aligned (0.699 ± 0.061). (C) HW fibers were significantly larger in diameter (1.61 ± 0.345 μm, p<0.01), and were less aligned (0.397 ± 0.076, p<0.01). (D) Tandem electrospun fibers of LW polymer solution and HW^+^ solution produced a centered pattern of fiber deposition with an overlapping gradient region between the two fibers also evident (inset). (E) LW had fiber diameters of 0.859 ± 0.311µm and were relatively aligned (0.551 ± 0.106). (F) HW^+^ fibers were smaller in size, though still significantly larger than LW fibers (0.865 ± 0.164 µm, p>0.01), and maintained the same approximate alignment as the original HW fibers (0.397 ± 0.076, p<0.01). (Scale bars: A,D 2 mm; Inset 500 μm)

Aiming to understand the sources of this bias, a comparison was made between the fluidic jet profiles with two identical LW solutions versus two different polymer solutions (Figure 1A Inset: “Tandem Jets” and Figure S4F, respectively). This revealed that T-ESP with identical polymer solutions produced stable jets of equal length while heterogeneous T-ESP exhibited different lengths, with the LW solution initiating the chaotic whipping phase at an earlier stage compared to the HW solution. This presented the possibility that the whipping phase of the LW jet electrostatically deflected the HW polymer jet to create the observed spatial bias.

To investigate this premise, the HW solution was replaced with the previously described HW*_mv_* solution (See Supplementary Table S2 for details). According to Hohman *et al.*^36^, the onset of whipping occurs once the polymer jet reaches a critical radius, with less viscous solutions reaching this stage earlier in the fluidic jet evolution. Despite the HW*_mv_* solution now initiating the whipping phase earlier compared to the LW solution, a spatially biased fiber pattern was still produced with LW still dominant on the target substrate (Supplementary Figure S5). Looking at the later stages of jet evolution showed that LW fibers experienced a final pull towards the target area (Supplementary Figure S5K). From this it was surmised that an LW fiber experienced a larger force during the final stages prior to deposition. This was attributed to the elongated whipping profile having more exposure to the electrostatic field and experiencing more electrostatic attraction.

To increase the effective force experienced by the HW fibers, the surface charge of the fibers was increased by adding 0.05 mg/ml of salt (NaCl) to the polymer solution^26^. As evidence of increased surface charge, this should also hasten the onset of the chaotic whipping phase^37^. NaCl was first added to the original HW solution (referred to hereafter as HW^+^), resulting in an increased conductivity (Supplementary Table S2). The increased surface charge of the HW^+^ fibers was evident from the whipping phase now initiating at approximately the same height as the LW jet (Supplementary Figure S6F). The end result was a more centered distribution of fibers (Figure 3E). The HW^+^ fibers were smaller, consistent with a previous report on the effect of salt^38^, and there was no change in alignment between the salt and salt-free condition.

Adding salt to the HW*_mv_* solution (hereafter HW*_mv_^+^*), evidence of increased surface charge was again observed by a much shorter stable jet length (approximately 1.5 cm) and a more centered distribution of tandem fiber deposition with a defined region of overlap (Supplementary Figure S7). The resulting HW*_mv_*^+^ fibers were now equal in diameter (0.774±0.287 μm) to the tandem-spun LW fibers (0.859±0.311 μm, p<0.01) but maintained a reduced degree of alignment (0.247±0.0586). In summary, tuning polymer solutions used for heterogeneous T-ESP makes possible fiber constructs with both distinct fiber alignments and tailored distribution patterns.

### Complex T-ESP fiber patterning

Once the optimization of T-ESP was established, the strategies for multi-jet patterned spinning were further extended. For example, moving from two to three jets resulted in the simultaneous overlapping deposition of three populations of aligned fibers (Figure 4A). The target electrode arrangement was further altered to achieve a divergent pattern of deposition (Figure 4C; Supplementary Figure S9 for electrode details). Figure 4B shows two fiber populations which have both deposited onto a ‘shared’ bottom electrode in an aligned, overlapping manner (Figure 4D). These fibers then diverge and separate, with the left polymer fibers (red) oriented towards the left upper electrode and the right polymer fibers (green) directed towards the right electrode. This creates a unique fiber pattern with a degree of complexity not yet seen for ESP.

**Figure 4.**
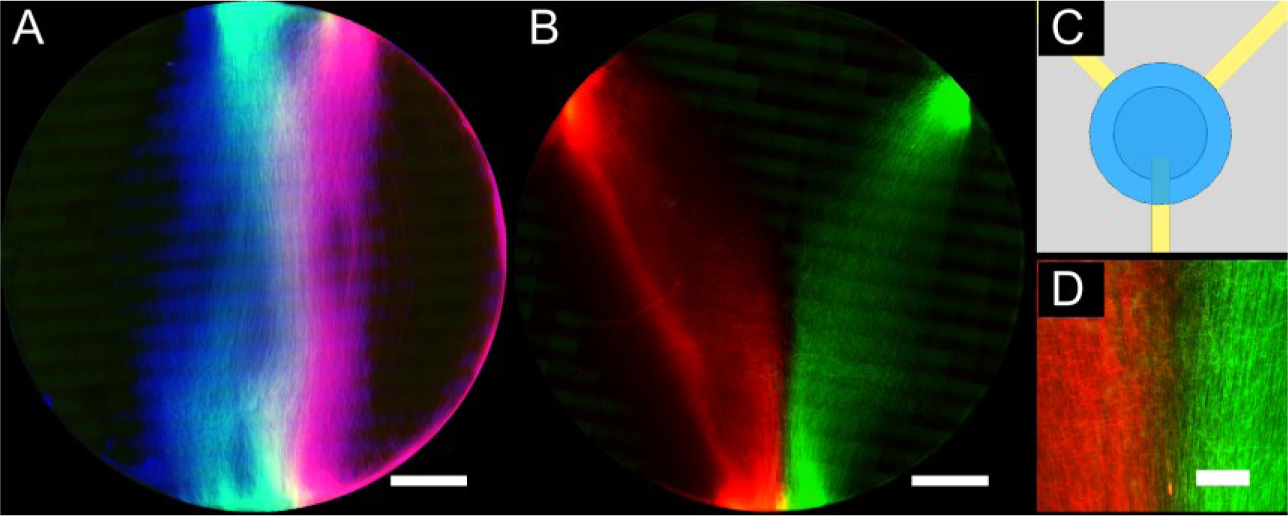
Triple ESP and Divergent Tandem ESP. (A) An example of focused aligned electrospinning with 3 separate LW solutions simultaneously (scale bar: 2 mm). A different dye was added to each solution (see Methods section). The blue autofluorescence of LW was enhanced by adding Pyrene, though the residual blue autofluorescence of the other fiber populations modified the typical green and red dye colors to mint and pink, respectively. (B) The result of tandem ESP on an ‘epsilon’ gap electrode arrangement (shown in C), producing a region of overlapping fibers (scale bars, B: 2mm; D: 500 μm) which then diverge. Further details of the needle and target collector arrangements for these patterns are described in Supplementary Material Figure S8. Fiber diameter is 0.788 ± 0.279 μm, consistent with previous LW ESP fibers.

### T-ESP applications

This new fabrication approach has an immediate application in the field of tissue engineering and regenerative medicine. The design of fibrous scaffolds follows a common biomimetic principle, with the intent of achieving an appropriate cell response by imitating the natural fibrous extracellular matrix found in the body. Current ESP scaffold can be tailored to elicit specific cell behavior, but fibers are typically modified homogeneously. However, this does not reflect the heterogeneous cell populations found *in vivo*. The possibility to create tissue scaffolds with purposely-designed heterogeneity may enhance scaffold effectiveness, exacting spatial control of cell response imparted by the properties of respective fiber types. Furthermore, bifurcated branching structures are ubiquitous throughout the body, including, but not limited to, nerves, vasculature, pulmonary and breast tissue^39–42^. The ability to make fibrous scaffolds as described in this work provides a promising tool for studying these types of biological systems as well as for the development of tissue engineering solutions.

To show the utility of T-ESP fiber tissue scaffolds, we developed a heterogeneous divergent scaffold of LW fibers and an LW-collagen blended fibers (Figure 5A) to explore neurite outgrowth of an explant dorsal root ganglion (DRG). An earlier study showed modulated neurite growth on ESP scaffolds of PCL/collagen blended fibers compared to scaffolds of PCL-only fibers^43^. We were able to replicate these earlier findings on our heterogeneous scaffold, with longer and more consolidated neurite extension on LW-collagen fibers compared to LW fibres (Figure 5C). Neurites also followed the fiber orientation, creating a divergent growth pattern and lending itself to quantification via radially segment Scholl analysis to provide quantification of differential outgrowth (Figure 5D-E). This presents interesting possibilities for neural tissue engineering, such as the spatially selective promotion of different neural subpopulations along divergently oriented fibers, or a general tissue engineering approach for sorting of a heterogeneous cell population.

**Figure 5.**
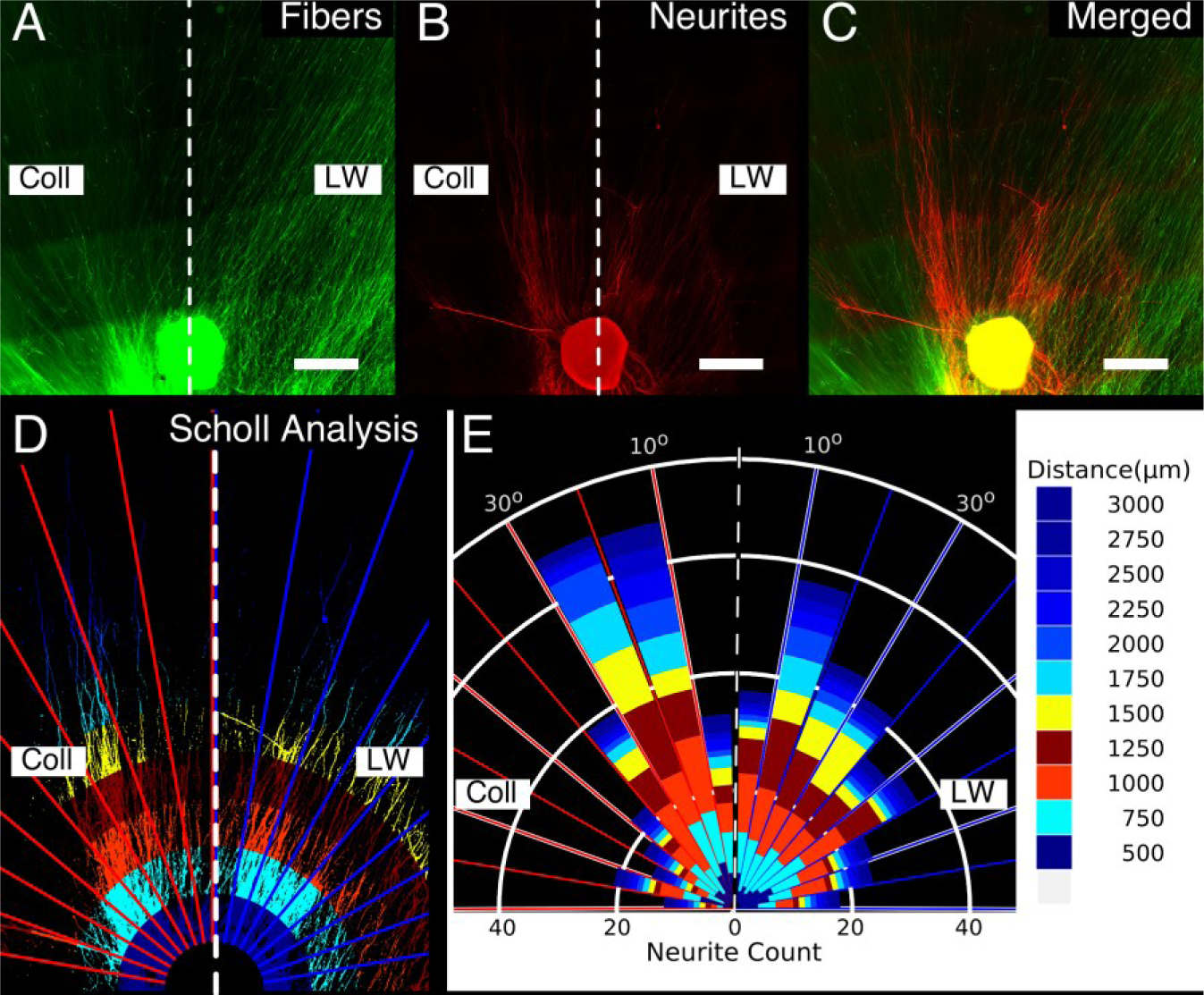
Spatially modulated neurite outgrowth on a divergent heterogeneous Tandem ESP scaffold. (A) A divergent heterogeneous T-ESP pattern comprised of LW-collagen fibers diverging to the left half labeled ‘Coll’ (0.235±0.048 m diameter; faintly autofluorescent green) and LW fibers diverging to the right half labeled ‘LW’ (with MacrolexGreen dye, visible as bright green; scale bar 500 μm). (B) Sensory neurite outgrowth from a rat Dorsal Root Ganglion placed at the junction of a divergent tandem ESP. Neurites were stained for 3-tubulin, shown in red (scale bar 500 μm). (C) A merged image shows neurite alignment along the direction of the divergent fibers and reveals a differential pattern of growth, with the collagen-containing fibers promoting longer, more consolidated growth while the LW fibers promote more shorter, dispersed neurites (scale bar 500 μm). (D) A visual representation the segmented Scholl analysis performed to measure neurite outgrowth. The image was divided into 10° angular sections, indicated by red lines for the LW-collagen half and blue for the LW half. Each sector was divided radially into 250 m segments indicated by the different colored bands. This created a 2D polar bin map with which the number of neurites per bin were then counted according to standard Scholl Analysis. (E) A rose plot of the cumulative number of neurites per bin. Each radial distance is indicated by the color and the size of each colored segment represents the neurite count within that distance. This analysis clearly shows that the LW-collagen fibers promote longer growth consolidated over fewer sectors compared to the LW fibers, which show dispersed growth over a broader range of sectors.

A gradient of fiber alignment could also spatially modulate cell morphology and, thus, creating a gradient of cell function. Figure S10 shows the modulation of Schwann cell morphology on a T-ESP scaffold of aligned LW fibers and less aligned HW*_mv_*^+^ fibers, where such differences in Schwann cell morphology are known to modulate their production of neurotrophic factors^44^.

The application of this technology is not limited to the field of Tissue Engineering. Initial trials have also employed tandem scaffolds to separate mixed solutions on the basis of fiber affinity (Supplementary Material S11), showing promise in the field of Separation Technologies. This fabrication technique can used to prepare heterogeneous catalytic or filtration substrates or high surface area nanofibrous cathode/anode electrode configurations able to sequester specific analytes.

## Conclusions

Until now, the ease and promising characteristics of ESP has found limited applicability due to limitations in the ability to control and customize the resulting fiber meshes. This work presents the highly accessible T-ESP approach, providing the versatility to create an array of micro- and nano-scale fibrous constructs with complex, ordered patterns. At the same time, this work provides general considerations for optimal polymer selection as well as furthering the understanding of the ESP process. Though a promising tool for tissue engineering applications, this approach holds potential for other fields. The methodology presented here further extends the growing arsenal of ESP solutions to create increasingly complex polymer fiber constructs in a simple manner to address the growing demands of new material and engineering challenges.

## Acknowledgements

The authors would like to thank Dr. Maqsood Ahmed, Dr. Alvaro Gomez Marin, and Dr. Hugo Fernandez for enlightening discussion during the experimental work and Mijke Buitinga for assistance in manuscript preparations. This work was partly funded by the National Science and Engineering Research Council (NSERC) of Canada and by funding received from the European Community’s Seventh Framework Programme (FP7/2007-2013) under MERIDIAN (grant agreement N°280778).

## Supplementary Information

### Experimental Section

#### Polymer solution preparation and characterization

All PEOT/PBT polymers were dissolved over night in solvent solutions of chloroform (CHCl_3_) and hexafluoroisopropanol (HFIP) in the ratios listed in Table S1. Macrolex® Fluorescent Yellow 10GN and Fluorescent Red G hydrophobic dyes (Lanxess NV, Belgium) were added at 0.01% w/v to assist in visualization. A 10mg/ml NaCl (Sigma Aldrich, Germany) suspension in HFIP was added to HW-based polymer solutions to final concentration of 0.05 mg/ml.

Conductivity was measured using a custom gold parallel plate apparatus. Applied voltage of approximately 0.707 V_rms_ was applied by a function generator(5400A Generator, Krohn-hite) at a frequency of 1 kHz. The voltage across the ‘sense’ resistor (R_sense_), connected in series with the probe, was used to determine the current through the circuit (Figure S9); R_sense_ was adjusted to match the voltage drop across the probe. The voltage was measured using a Tenma 72-7725 MultiMeter. The conductivity probe was calibrated with a solution of 1 mol/L NaCl in MilliQ water with a known conductivity of 85 mS/cm. Viscosity was measured using a Brookfield DV-E with an s21 mandrel at 30 °C and 100 rpm.

For additional electrospun patterns (SI Appendix, Figure S8), 5% w/v solutions of Polyethylene Oxide (PEO, 900,000 MW, Sigma Aldrich) were prepared in demineralized water. To distinguish different PEO fiber populations, solutions were prepared with Fluorescein (green, 0.01% w/v, Sigma Aldrich), Rhodamine (red,0.1% w/v,) and 4′,6-Diamidino-2-phenylindole dihydrochloride (DAPI, blue, 0.01% w/v, Sigma Aldrich). The PEOT/PBT-collagen blend was prepared by first preparing a 20% w/v LW solution and an 8% w/v collagen in HFIP. These were then mixed in a 1:1 ratio for a final solution of 10% w/v LW and 4% w/v collagen. A 10% solution of polyvinyl alcohol (PVA, 98% anhydrous, Sigma Aldrich) was prepared in demineralized H_2_O. A mixed drop of corn oil with hydrophobic MacrolexGreen (20 □g/ml) and water with hydrophilic Rhodamine B (20 □g/ml) is placed on the central region of the scaffold. A time series was captured using a non-inverted Nikon E600 and a triple band pass filter (DAPI/Green/Red).

#### Electrospinning

Electrospinning was performed in an environmental chamber, with a maintained temperature of 25 °C and a relative humidity from 22% to 27%. Polymer solutions were loaded into 5 ml syringes (BD Biosciences) and connected via Teflon tubing to a stainless steel needle (0.8 mm outer diameter, 0.5 mm inner diameter). Needles were mounted to the centre of a large upper parallel plate (30 cm by 20 cm) and centered over the collector target at a working distance of 20 cm. A voltage of 25 KV was applied to the upper parallel plate and needles (Gamma HV Research). For tandem spinning, needles were mounted along the Y axis of the collector target (Figure 1B) at an inter-needle separation of 1 cm; needles were symmetrically offset with respect to the X axis. Triple spinning used an inter-needle separation of 5 cm along the Y axis, with one needle centered at the YX axis intersection. Using a syringe pump (KDS-100-CE, KD Scientific), one per syringe, solutions were electrospun at flow rate of 1 ml/hr, unless otherwise stated. A 14 mm coverslip (Menzel-Glaser) was the target substrate used to collect deposited fibers, placed on the collector electrodes of either a parallel or epsilon arrangement (see SI Appendix, Figure S8).

#### Whipping and Fiber Analysis

Images of whipping were captured by a Luminix DMC G3 with bar LED illumination. Fiber patterns were recorded with a Nikon Eclipse Ti with an automated stage at 10x magnification in brightfield and with an epifluorescent lamp with filters for FITC, Rhodamine, DAPI and brightfield. Images were stitched using Nikon Elements software. Fibers were gold sputter-coated with a Polaron E5600 sputter-coater and examined with a XL 30 ESEM-FEG (Phillips) operating at 10 kV. Fiber diameter was determined by measuring a minimum of 100 fibers, taken from a minimum of 5 images at random locations with a 1000x magnification. A minimum of 5 images per fiber population were evaluated for fiber orientation using the OrientationJ plugin(1), providing a coherence value between 0 (isotropic) and 1 (perfectly anisotropic). Fiber orientation was also evaluated by creating an FFT image of SEM images (2). Images of whipping fibers, fiber patterns and fibers were processed with Fiji image processing software (3). Figures were prepared with Scientifig plugin (4).

#### Statistics

All statistics were performed with R statistical software (http://www.R-project.org/) and graphs were created using the Deducer plugin (by Ian Fellows, http://www.jstatsoft.org/v49/i08/). When comparing between tandem ESP fiber populations, a Student’s T test was used. When comparing whole populations of the same fiber family, a one way ANOVA was use followed by a *post hoc* Holm method. A significance level of p < 0.01 was adopted.

**Figure S1.**
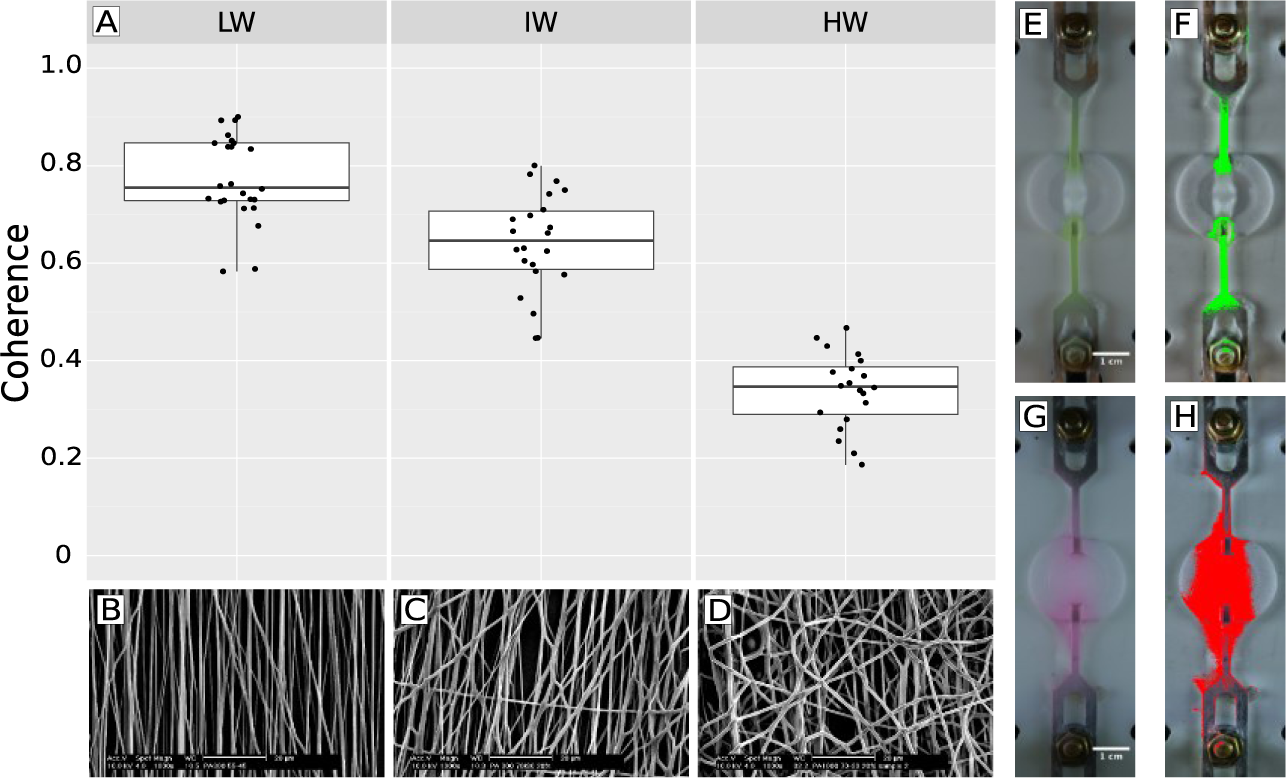
Fiber alignment of LW, IW and HW polymer solutions when electrospun individually (solo ESP versus tandem ESP). (A) plots fiber alignment, based on the Coherence metric provided by the OrientationJ plugin for ImageJ/Fiji (1). (B-D) show representative SEM images of the LW, IW and HW fibers. (E) shows LW, with MacrolexGreen dye, having an elongated deposition pattern along the electrodes, which becomes more apparent when applying a color thresholded to highly green pixels. In contrast, G shows a less elongated, more circular deposition of HW polymer fibers, with a similar threshold filter making this more clear in (H). (scalebar: E, G 1 cm).

**Figure S2.**
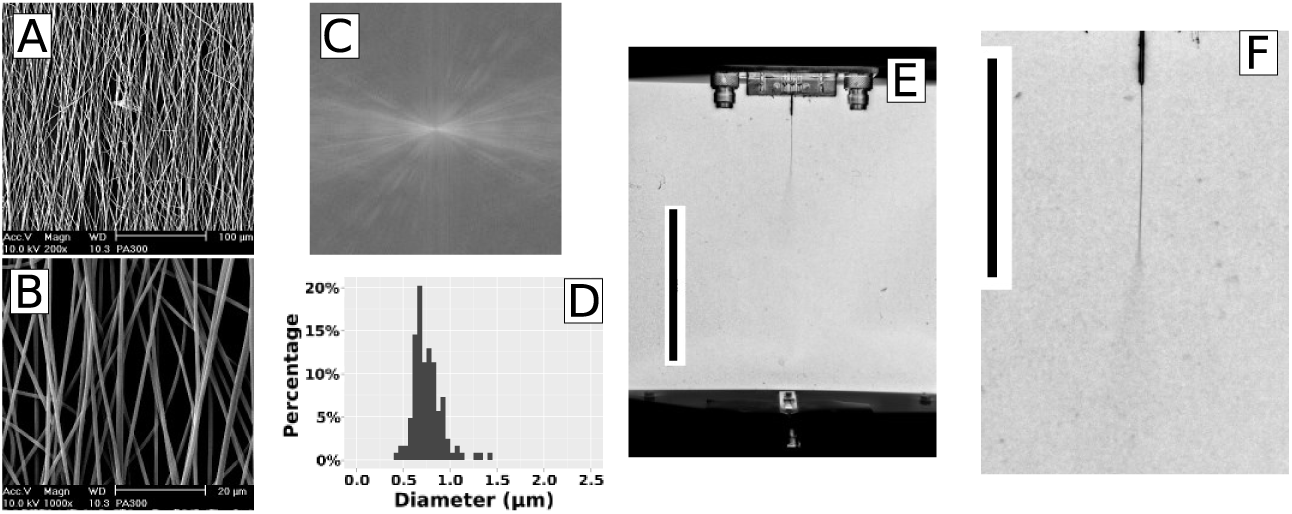
(A,B) SEM analysis of aligned ESP fibers of a single jet of LW polymer solution show they are well formed and aligned. Alignment is corroborated by indications of alignment in an FFT image of the SE micrograph (C) and an image coherence measurement of 0.780 ± 0.059 (1 is perfectly aligned, 0 is isotropic). Fibers had a diameter of 0.756 ± 0.160 μm, with a size distribution shown in (D). The whipping profile of a single jet is also shown in (E, scalebar: 10 cm), with a close up in (F, scalebar: 5 cm) showing a stable jet length (thin dark line) of approximately 3.5 cm.

**Figure S3.**
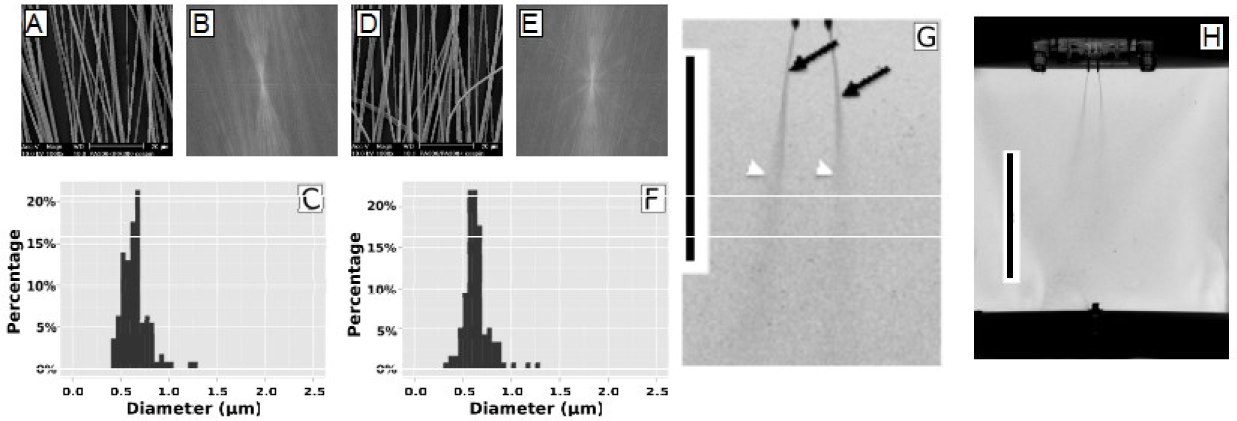
SEM images of tandem ESP of two LW solutions on a 14 mm coverslip (A,D) show parallel depositions of aligned fibers of the two fibre populations with similar degrees of fiber alignment (0.734±0.115 and 0.817±0.097, FFT of SEM images in (B) and (E)). Both fiber populations were of similar diameters (0.653±0.143 μm; 0.143±0.136 μm) and had similar fiber diameter distributions (C, F). Whipping jet profiles (scale bars: G 5 cm; H 10 cm) shows equal stable jet lengths and initiation of whipping at the same point.

**Figure S4.**
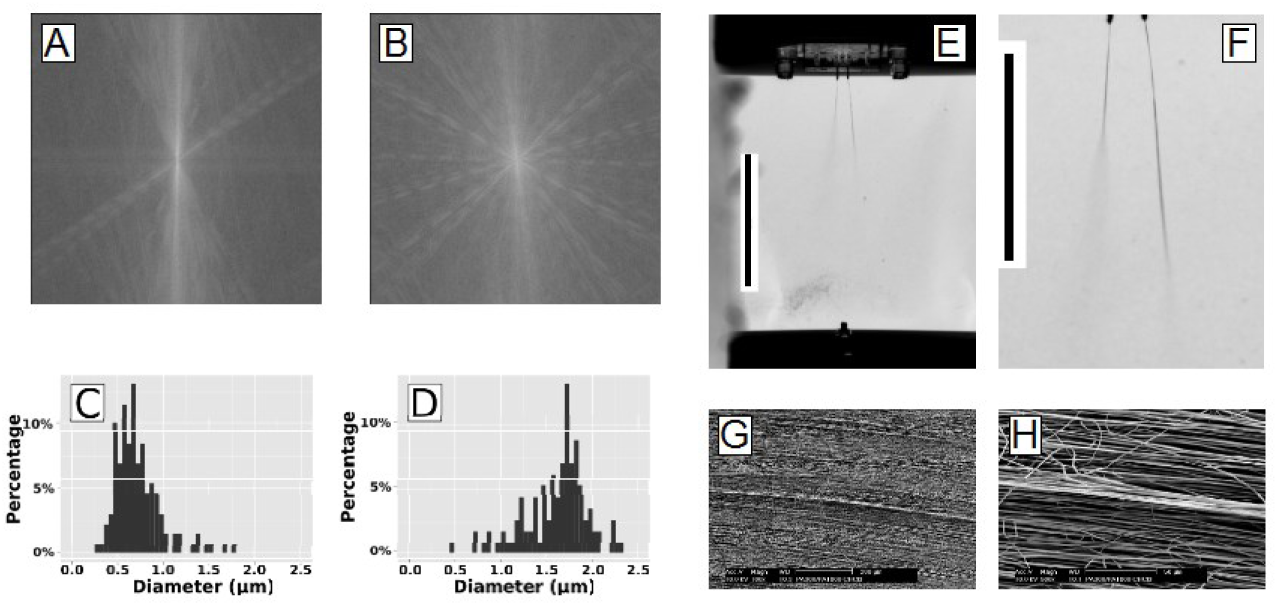
Tandem ESP of LW and HW polymer solutions produce LW fibers with a coherence of 0.699 ± 0.061, indicating fair alignment (FTT shown in (A), and a diameter of 0.723 ± 0.261 μm, distribution shown in (C). In comparison, the HW fibers were less aligned (0.397 ± 0.076, FFT (B), and had a diameter of 1.61 ± 0.345 μm (histogram shown in (D)). The whipping jet profiles (E, scale bar: 10 cm; F, scale bar: 5 cm) reveal that the HW solution initiates whipping later compare to the LW solution. Distinct ridges of accumulated parallel fibers are noted at the transition between the two fiber types (G, H), a possible consequence of electrostatic interactions between the two fiber populations.

**Figure S5.**
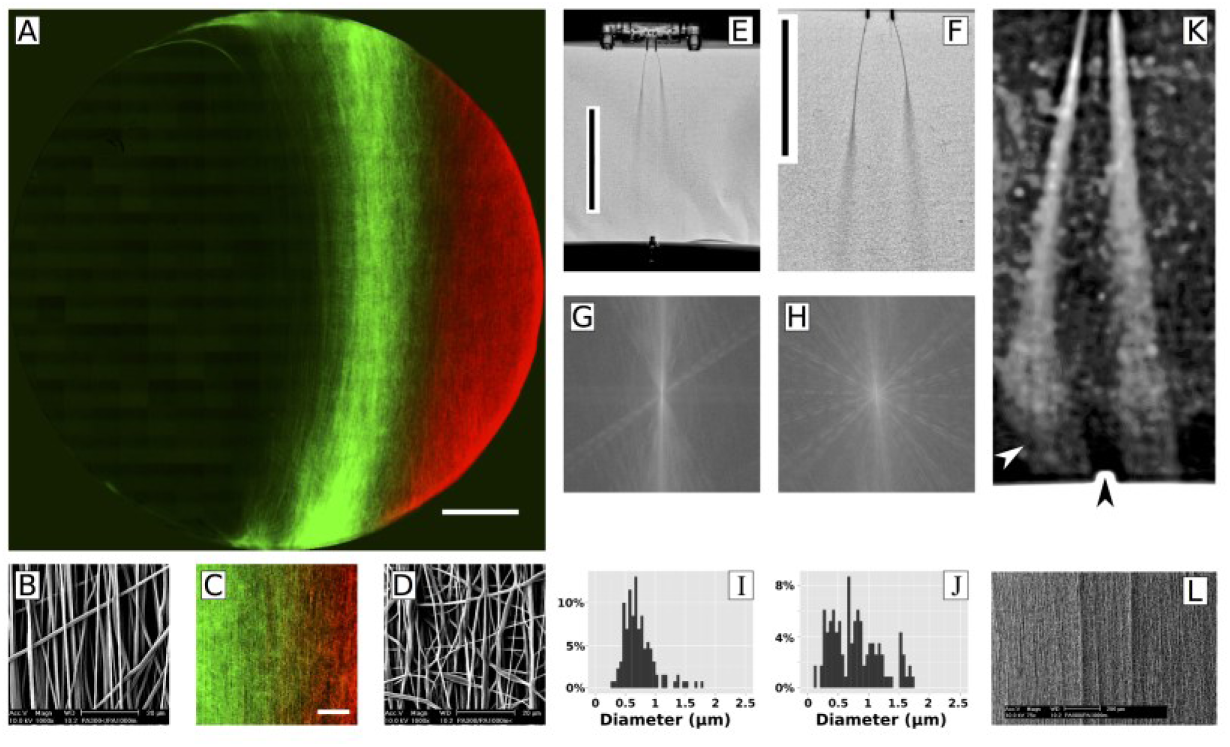
Tandem ESP of LW and HW_mv_, which produces a biased deposition of fibers (A, scale bar: 2 mm) as well as a region of overlap (C, scale bar: 500 μm). Looking at the stable jet phases of the tandem ESP (E, scale bar: 10 cm; F, scale bar: 5 cm), the lower viscous HW_mv_ had a shorter stable jet length (right) compared to the LW jet (left) An image of the lower whipping phase (K), processed with Fiji to increase contrast (imaged was processed with functions: smooth, CLAHE local contrast enhancement, background substraction) to make the small fibers visible, shows the LW jet is deflected by HW_mv_ whipping but experiences a final ‘pull’ of the LW fibers (left jet, white arrow) towards the electrode area (indicated by the black arrow, center bottom). In comparison, the right HW_mv_ jet shows no such redirection. Based on this observation, the assumption was made that the biased deposition was the result of greater electrostatic force upon the LW fibers at the final stages of deposition. The transition region again produces accumulated bundles of fibers, forming striated like structures (L). LW fibers are of similar diameter to previous LW fibers (0.647±0.223 μm) and share a similar distribution (I). As well, alignment is also similar (0.804±0.049, (G)). The fibers of the less viscous HW_mv_ solution produce a smaller fiber diameter (0.790 ± 0.402 μm) than the previous HW solution and have a slightly improved alignment, though not statistically significant (0.432 ± 0.076).

**Figure S6.**
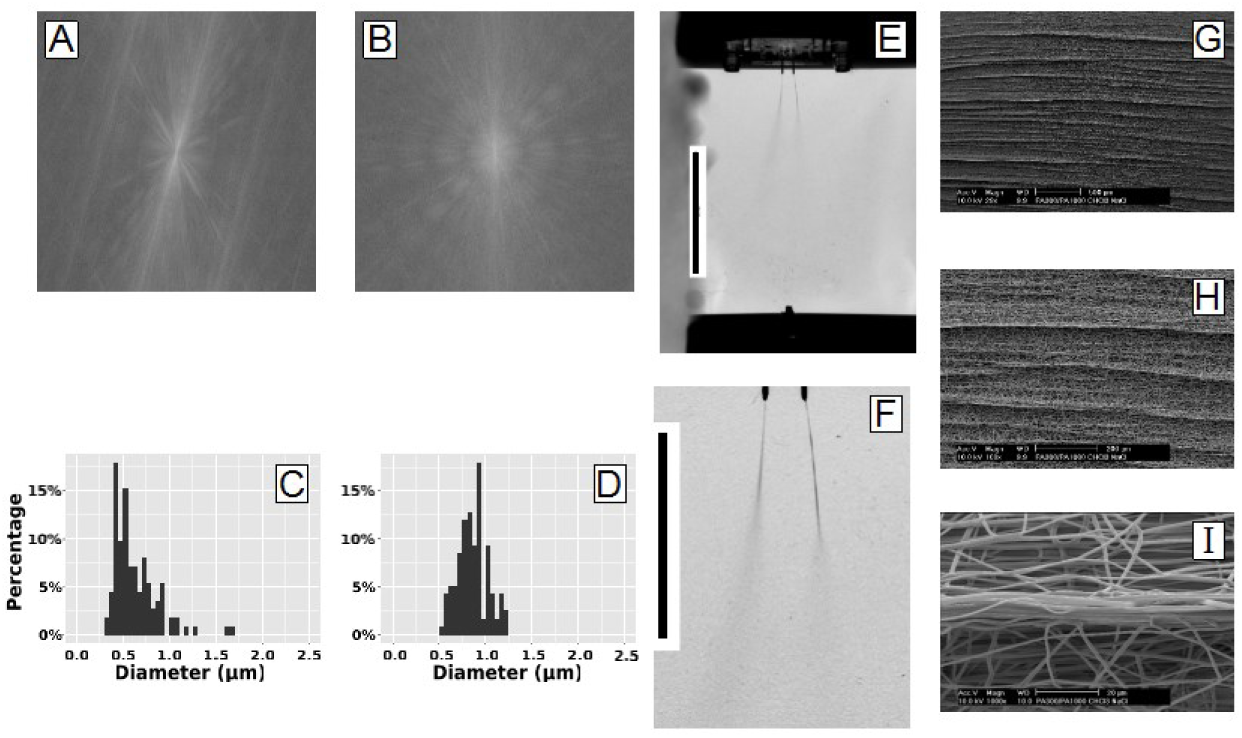
Tandem ESP of LW and HW^+^ polymer solutions produces LW fibers with an alignment of 0.633±0.061 (corresponding FFT image (A)) and a diameter of 0.630 ± 0.242 μm, with the diameter distribution shown in (C)). The HW^+^ fibers have a smaller diameter compared to the original HW solution (0.865±0.164 μm) and a narrower diameter distribution (D), though alignment is roughly equal (0.397±0.076 (corresponding FFT image (B)). The whipping profiles shown in (E) and (F) show that the LW jet (left) and the HW^+^ jet have similar stable jet lengths (scale bars of 10 cm and 5 cm, respectively). As before, the transition region exhibits striated patterns of highly aligned fiber bundles (G-I).

**Figure S7.**
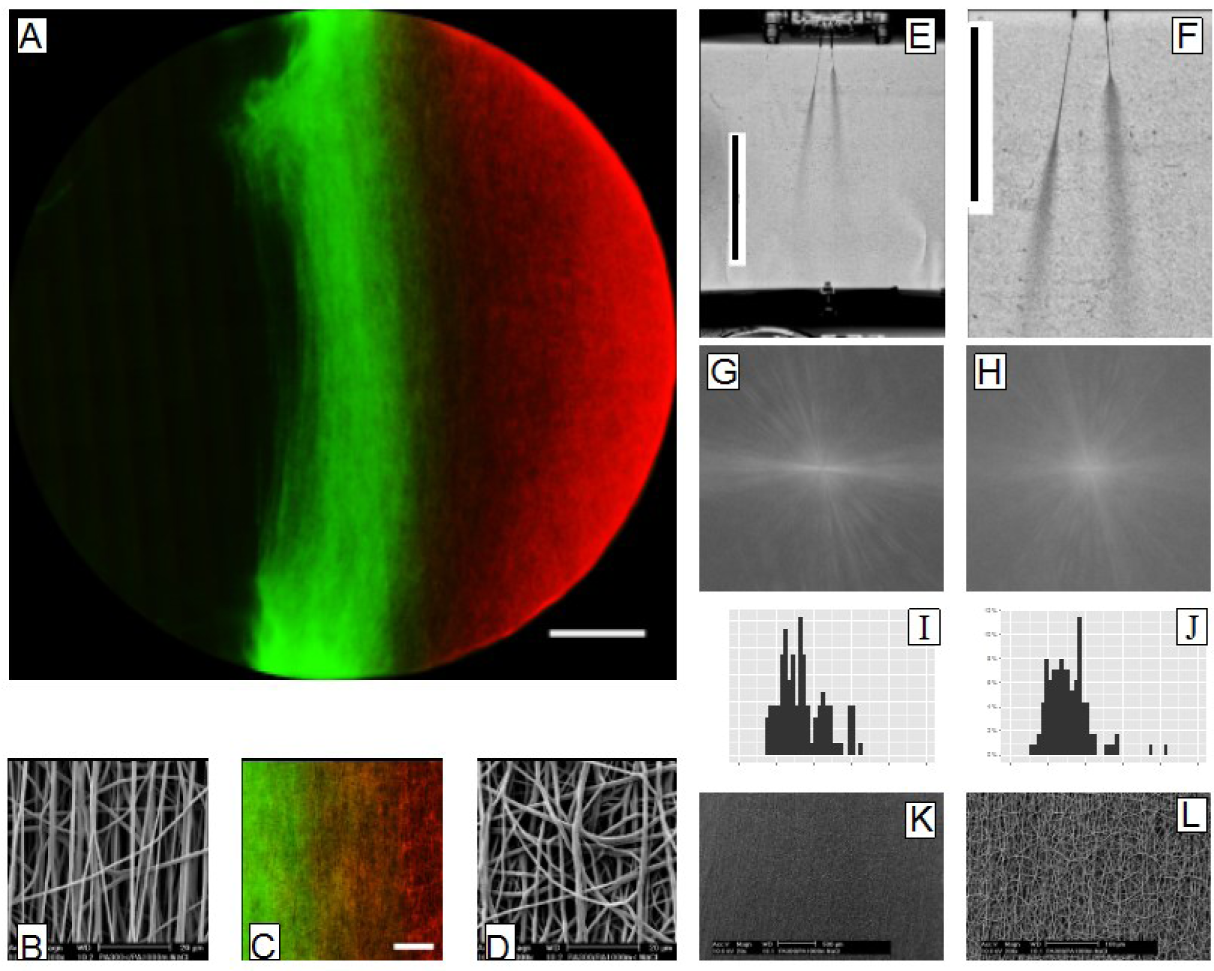
Tandem ESP of LW and HW_mv_^+^ solutions produced a centered fiber distribution (A, scale bar: 2 mm), this time with a region of overlap (C, scale bar 500 μm) which did not result in a striated transition region (K,L). The jet profiles show that the HW_mv_^+^ stable jet (left) is now much shorter than the LW stable jet (right) (E, scale bar: 10 cm; F, scale bar: 5 cm). LW fibers had a diameter of 0.859 ± 0.311 μm, larger than previous LW fibers and with a wider distribution (I), and have comparatively less alignment (0.551 ± 0.106, SEM image (B) and corresponding FFT image (G)). HW_mv_^+^ fibers have a diameter of 0.774 ± 0.287 μm (diameter distribution shown in (J)), equal to the tandem LW fibers produce, and exhibit a low degree of alignment of 0.247 ± 0.059, corroborated by SEM (D) and FFT analysis(H).

**Figure S8.**
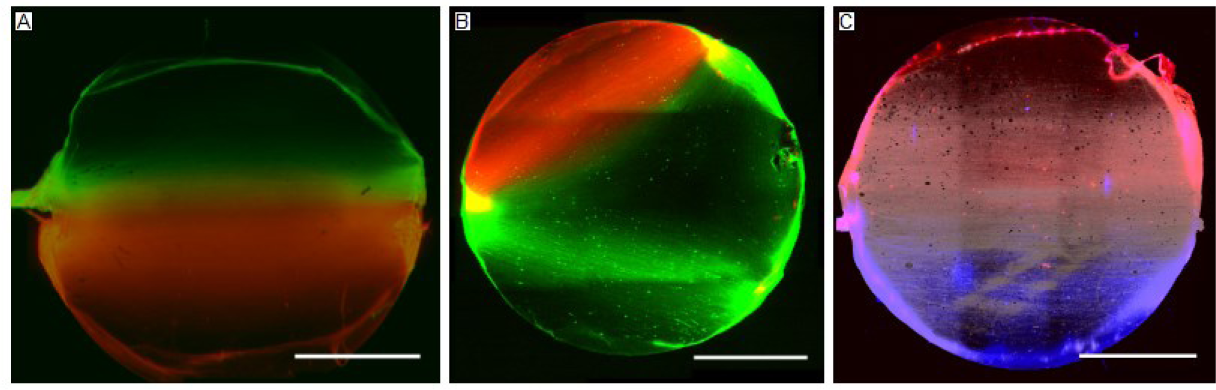
Additional examples of patterned spinning with a 5% PEO solution (900,000 MW). (A) Using the parallel electrode arrangment, a similar pattern can be achieved with this polymer solution. (B) A divergent pattern can also be achieved, though completely separation of polymer types was less roboust and subject to fine adjustment in needle alignment. Note the presence of green fibers diverging in the same direction as red fibers. (C) As well, a triple spin is possible with PEO. The central PEO fibers, with fluorescein, were adjusted to false white to improve visualization. (scalebars: 4 mm)

**Figure S9.**
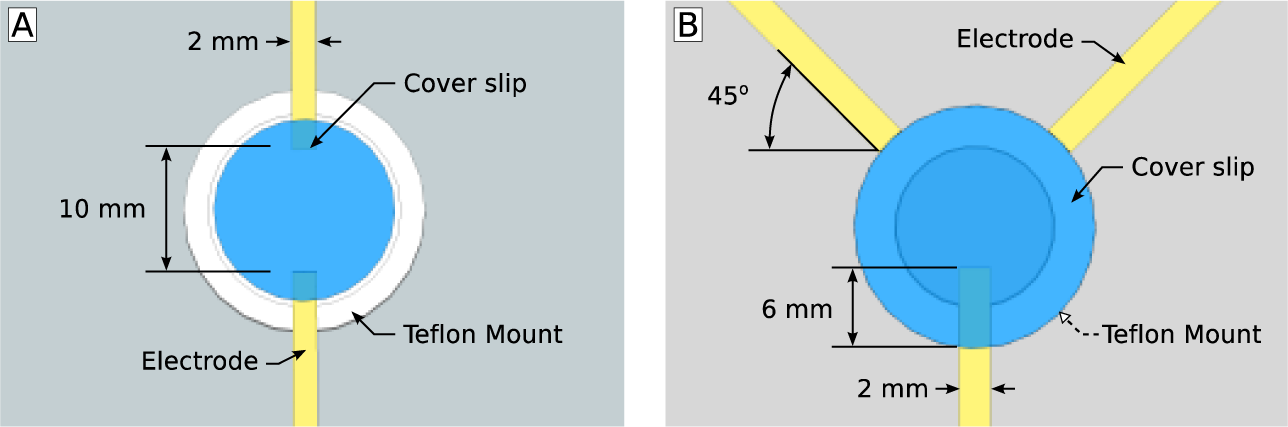
(A) The electrode arrangement used for parallel tandem ESP, with a gap width of 10 mm and electrode width of 2 mm. A Teflon mount was used to center a 14 mm coverslip (Menzel-Glaser) between the electrodes with a 2 mm of overlapping coverlsip over each electrode. The Teflon mount raised the cover slip 2 mm above the electrodes to ensure no direct contact. Tandem ESP with two jets used an inter-needle separation of 1 cm, centered about the coverslip along the X axis. Triple ESP used an inter-needle separation of 5 cm to ensure jet stability. (B) The divergent patterns were created using an epsilon-type gap electrode triple electrode arrangement shown. The base electrode repositioned 6 mm under the glass coverslip while two additional electrodes where positioned at the cover slip periphery at an angle of 45° with respect to the coverslip center. As before, an inter-needle separation of 1 cm, centered about the coverslip along the X axis. The coverslip was again placed on top of a Teflon mount to maintain position and ensure no direct contact with electrodes.

**Figure S10.**
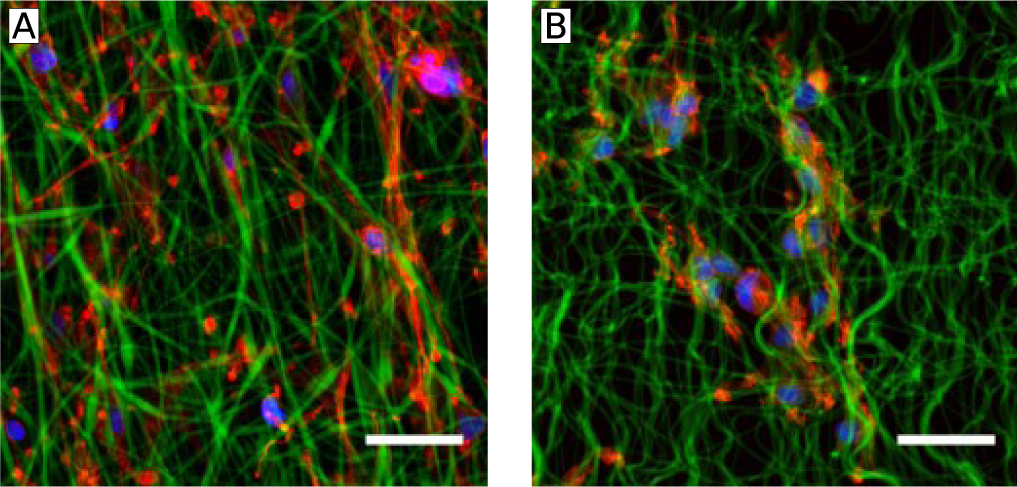
Schwann cells on a heterogeneous LW/HW Tandem ESP scaffold. Rat Schwann cells cultured on a tandem ESP scaffold of CA (strongly cell adherent) and NA (less cell adherent) polymer fibres. (A) Cells on CA show clear cell adhesion and elongation in the direction of fiber orientation, while (B) shows that cells adhere poorly to NA fibres and retain a rounded, less elongated cell morphology. Also notable is the wave-like morphology of the NA fibres, the result of swelling because of the high hydrophilicity of this polymer (scale bar: 20 μm).

**Figure S11.**
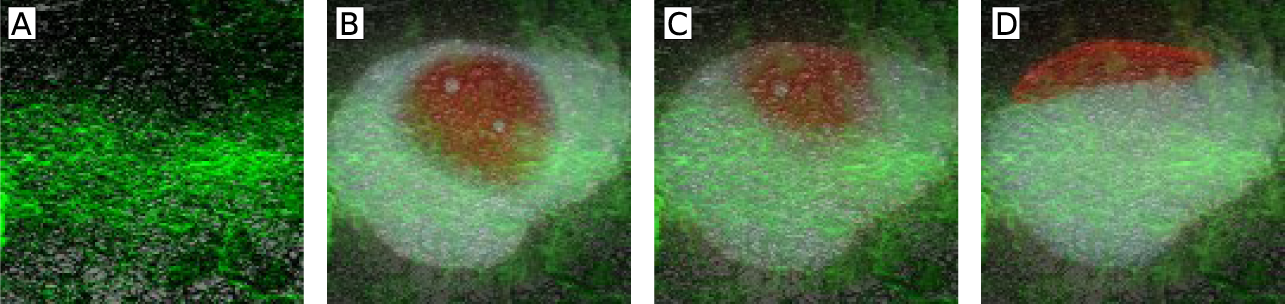
Oil/Water separation on a tandem ESP scaffold of LW (hydrophobic) and PVA (hydrophilic) fibers. (A) A merged image showing the location of LW (green) and PVA (black) fibers. A mixed drop of corn oil with hydrophobic MacrolexGreen (20 g/ml; shown as white) and water with hydrophilic Rhodamine B (20 □g/ml) is placed on the central region of the scaffold and imaged after (B) 30 seconds, (C) 2 minutes and (D) 12 minutes. Observed is migration and separation of the two phases in accordance with the underlying fibers.

**Figure S12.**
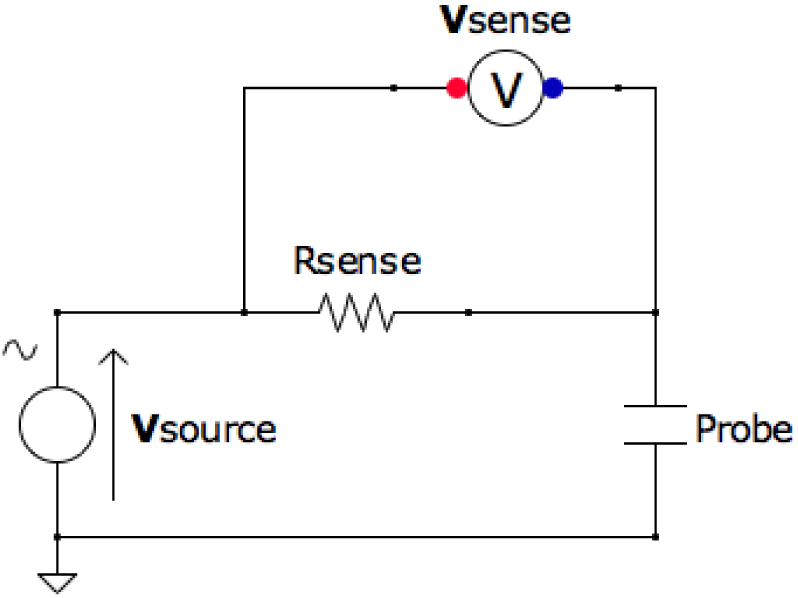
A depiction of the circuit used to measure the conductivity of polymer solutions. By measuring the V_sense_ across the known R_sense_, i_sense_ could be calculated. The voltage across the probe (V_probe_) was calculated by substracting V_sense_ from V_source_ By applying Ohm’s Law to i_sense_ and V_probe_, the resistance of the polymer solution is determined. Conductivity was calculated using the empirically derived constant of the probe using a dilution series of Glycerol with NaCl concentrations and compared against published values of conductivity.(5)

**Table S1.** Polymer Properties^a^

| Polymer | Mw<br>(kDa)<sup>a</sup> | Me<br>(Da)<sup>a</sup> | H<sub>a</sub><br>(MPa)<sup>a,b</sup> | PEO segment<sup>c</sup> |  | PBT segment<sup>c</sup> |  |
| --- | --- | --- | --- | --- | --- | --- | --- |
|  |  |  |  | Tg<br>(°C) | Tm<br>(°C) | Tg<br>(°C) | Tm<br>(°C) |
| 300PEOT55<br>PBT45 | 50.67 | 250 | 97.15 | -23 | -<sup>d</sup> | 26 | 145 |
| 300PEOT70<br>PBT30 | 48.45 | 450 | 42.7 | -23 | - | - | 114 |
| 1000PEOT7<br>0PPBT30 | 57.43 | 710 | 29.6 | -50 | 6 | - | 149 |

<sup>a</sup> Data taken from Moroni et al., 2005(6)<sup>b</sup> H<sub>a</sub> is the equilibrium modulus<sup>c</sup> Data take from Deschamps et al., 2001(7)<sup>d</sup> not observed**Table S2. Polymer Solution Properties**

**Table S2.**
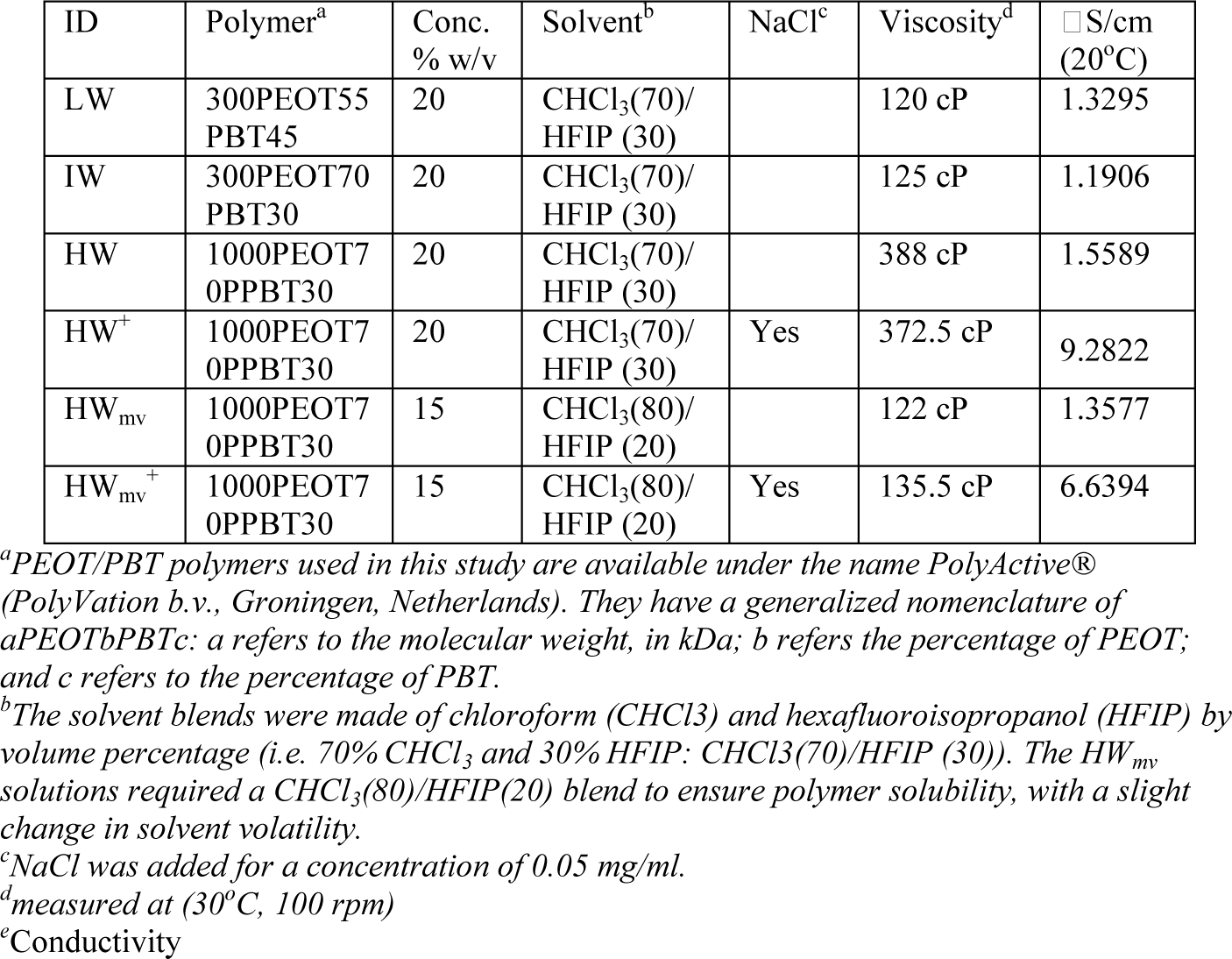
Polymer Solution Properties

**Table S3.**
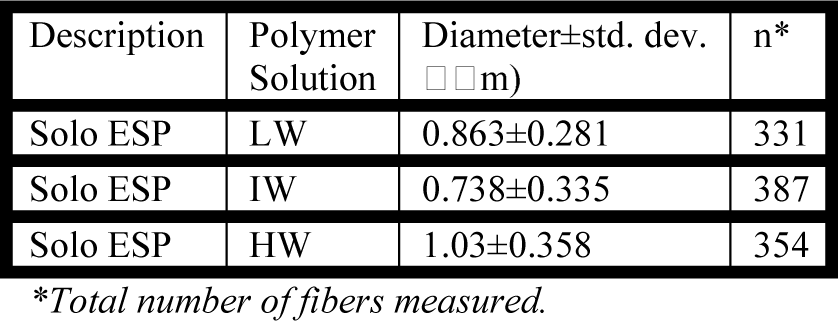
Summary of ‘solo’ ESP Fiber Diameters

**Table S4.** Summary of Tandem ESP Fiber Diameters

| Description | Polymer Solution | Diameter±std. dev. (μm) | n* |
| --- | --- | --- | --- |
| Tandem ESP | HW<sub>mv</sub> | 0.790±0.402 | 115 |
| | LW | 0.647±0.223 | 149 |
| Tandem ESP | HW<sub>mv</sub><sup>+</sup> | 0.774±0.287 | 113 |
| | LW | 0.859±0.311 | 107 |
| Tandem ESP | HW | 1.607±0.345 | 113 |
| | LW | 0.723±0.261 | 130 |
| Tandem ESP | HW<sup>+</sup> | 0.865±0.164 | 117 |
| | LW | 0.630±0.242 | 112 |
| Tandem ESP | LW | 0.653±0.143 | 108 |
| | LW | 0.637±0.136 | 118 |

\*Total number of fibers measured.

**Table S5.** Summary of Solo ESP fiber alignments

| Description | Polymer Solution | Coherence±std. dev.* | n** |
| --- | --- | --- | --- |
| Gap Electrode | LW | 0.773±0.089 | 24 |
| Gap Electrode | IW | 0.641±0.102 | 22 |
| Gap Electrode | HW | 0.339±0.077 | 20 |
| Random ESP | LW | 0.088±0.055 | 7 |

\*Coherence as a metric for orientation of objects in an image, as measured by the OrientationJ plugin(1). 1 is perfect alignment while 0 is complete anisotropy.
\*\*Total number of images analyzed.

**Table S6.** Summary of Tandem ESP fiber alignments

| Description | Polymer Solution | Coherence±std. dev.* | n** |
| --- | --- | --- | --- |
| Tandem ESP | HW<sub>mv</sub> | 0.432±0.076 | 6 |
| | LW | 0.804±0.049 | 7 |
| Tandem ESP | HW<sub>mv</sub><sup>+</sup> | 0.247±0.059 | 6 |
| | LW | 0.551±0.106 | 6 |
| Tandem ESP | HW | 0.401±0.060 | 7 |
| | LW | 0.699±0.104 | 6 |
| Tandem ESP | HW<sup>+</sup> | 0.397±0.076 | 7 |
| | LW | 0.633±0.061 | 6 |
| Tandem ESP | LW | 0.734±0.115 | 6 |
| | LW | 0.817±0.097 | 6 |

\*Coherence as a metric for orientation of objects in an image, as measured by the OrientationJ plugin(1). 1 is perfect alignment while 0 is complete anisotropy.
\*\*Total number of images analyzed.

